# High Sensitivity of *Tropilaelaps mercedesae* to Lithium Chloride: A Novel Acaricidal Candidate for Honey Bee Health

**DOI:** 10.64898/2025.12.15.694363

**Authors:** Éva Kolics, Pichet Praphawilai, Bajaree Chuttong, Judit Poór, Balázs Kolics

## Abstract

The escalating threat of the ectoparasitic mite *Tropilaelaps mercedesae* requires novel control strategies for honey bee (*Apis mellifera*) health^4^. In this study, we provide the first evidence of the acaricidal efficacy of lithium chloride (LiCl) against this emerging parasite. *Ex situ* contact bioassays quantified the dose-response relationship, revealing a 12-hour *LC*_*50*_ of 45.9 mM. These results indicate that *T. mercedesae* is intrinsically more than 3.5 times more sensitive to LiCl than the widespread *Varroa destructor*. We translated these findings to field conditions in a pilot trickling trial on infested *A. m. ligustica* colonies. Repeated administration of 500 mM lithiated sugar syrup induced characteristic tremors and a sharp increase in mite fall, confirming that the acaricidal effect manifests *in situ* even in the presence of brood. Our findings identify lithium as a highly effective, naturally occurring candidate for *Tropilaelaps* management. This warrants larger-scale trials to refine application protocols and evaluate long-term colony safety for sustainable apiculture.

## Introduction

The western honey bee, *Apis mellifera*, is a crucial insect pollinator, and its managed populations are fundamental to global food security. Commercial pollination services are overwhelmingly dependent on this single species, which underpins a significant portion of the economic value derived from agricultural pollination. Consequently, the introduction of invasive species through intense global trade poses a serious, often underestimated, threat to this vital ecosystem service ^1^. Currently, the primary biotic driver of honey bee colony losses worldwide is the ectoparasitic mite, *Varroa destructor*, which has become nearly ubiquitous in all regions where *A. mellifera* is kept ^2^.

However, an equally, if not more, pernicious parasite, the *Tropilaelaps* mite, particularly *T. mercedesae*, represents an escalating threat. Having 20 haplotypes found in its indigenous host *A. dorsata*, 6 haplotypes have been found on *A. mellifera* ^3^, suggesting this mite is undergoing a successful and continuous expansion from its native range in Asia, increasing the risk of its introduction to more distant continents.

The current distribution of *T. mercedesae* primarily encompasses the Indochinese region including Thailand, Myanmar, Nepal, and China, where it is prevalent alongside its indigenous host, *Apis dorsata*. However, it has also successfully spread within *A. mellifera* colonies in the Philippines, however, no infestation has been detected in *A. mellifera* colonies on the Palawan Islands ^3^. Conversely, in South Korea, *T. mercedesae* populations are established ^4^ and have shown continuous growth since their first appearance (Jung et al., 2014). Notably, the successful establishment of this mite in this temperate climate, where winter can be harsh and honey bee brood rearing limited, suggests *T. mercedesae*’s ability to adapt and thrive in colder environments, similar to *Varroa* mites. Moreover, recent research has shown that *T. mercedesae* can exit colonies by phoresy on adult *A. mellifera*, indicating a potential for natural dispersal between colonies, especially when brood infestation is elevated ^5^.

Looking westward, while *Tropilaelaps clareae* (likely *T. mercedesae* based on Anderson and Morgan’s 2007 reclassification) was detected in Kenya, no established mite population has been reported there. Its presence in Iran was sporadicly reported ^6^ but its establishment in Iran remains doubtful ^7^. Recently, it has also made its way to Europe, first establishing in Russia ^8^, with the most recent detection occurring in Georgia ^9^. This latest expansion into the Caucasus region raises serious concerns, as the mechanism facilitating its persistence in colder climates remains unclear. Key unresolved questions include its ability to survive (i) short, non-broodless winter periods, (ii) a potential facultative diapause, or (iii) overwintering on an unknown alternative host. The rapid population growth of *Tropilaelaps* mites with worker brood infestation increasing from 0.40% to 15.25% in just 60 days in untreated colonies, leading to significant colony damage—underscores the urgent need for effective management ^5^. Its faster population growth of *Tropilaelaps* mites compared to *Varroa* - primarily due to a life cycle almost entirely confined to capped brood and a much shorter phoretic phase - suggests this parasite could pose an even greater threat than its infamous relative ^3^.

Controlling *Tropilaelaps* infestations presents a formidable challenge. While many conventional acaricides, such as amitraz or hops beta acids, are largely ineffective against *T. mercedesae*, formic acid has shown consistent efficacy due to its unique ability to penetrate cell cappings and target mites within the brood ^10^. However, the increasing emergence of resistance to synthetic acaricides is a growing concern. Previous studies have suggested that *Tropilaelaps* may exhibit resistance to active ingredients such as Amitraz, Coumaphos, Flumethrin, and Fluvalinate (Gill et al., 2024 ). Essential oils have demonstrated acaricidal efficacy against *Tropilaelaps* under laboratory conditions, with Piper betle showing the highest activity and several essential oils exhibiting low toxicity to adult honey bees. However, their efficacy in colony conditions requires further investigation regarding optimal concentrations and delivery methods ^11^. Exploiting the mite’s brood dependency, beekeeper-induced brood breaks have proven to be a highly effective cultural control method, even without chemical intervention. However, the combination of a brood break with a single chemical treatment (formic acid or oxalic acid), while significantly reducing mite populations, has not achieved complete eradication, necessitating further optimization for eradication efforts ^10^.

Given the potential threat of resistance development observed in *Tropilaelaps* mites, similar to that experienced with *Varroa destructor*, there is an urgent need for research and development of new and effective treatment strategies. This, however, presents a formidable challenge. An ideal active substance must be less toxic to bees than the target organism, not prone to accumulating in water-soluble or lipophilous bee products like honey and beeswax, and be administrable with high efficacy to eradicate the pest, which is difficult to access as it propagates within the capped brood.

Recently, a novel potential active ingredient, lithium, has emerged. Its acaricidal properties were first uncovered in *V. destructor* ^12^. This study was prompted by an observation from an earlier experiment on siRNA-based gene silencing ^13^, where lithium chloride, used merely as a stabilizing agent for the dsRNA in the control group, was unexpectedly revealed to have potent acaricidal effects. Lithium shows promise due to both its systemic ^12^ and contact action ^14^. Crucially, even after high exposure, residues have been found to be negligible in wax matrices and do not accumulate to alarming levels in honey ^15, 16^. Furthermore, lithium is a natural trace element in some floral honey types ^17, 18^. While further validation is required for regulatory approval, its practical relevance is supported by studies demonstrating its efficacy also in comparison to alternatives which can also be considered as a natural component of honey (eg. oxalic acid ^19^), via application methods, such as trickling, that are congruent with current beekeeping practices for acaricide treatments ^20^.

Recent investigations have revealed that the acaricidal activity of lithium extends beyond *Varroa* to other taxonomically distant mite and tick species, including the tick *Dermacentor reticulatus* ^21^, the poultry red mite *Dermanyssus gallinae* ^22^, and the two-spotted spider mite *Tetranychus urticae* ^23^.

The objective of our study was therefore to determine whether the efficacy of lithium extends to *Tropilaelaps mercedesae* and, if so, to evaluate its potential for practical application in apiculture.

## Materials and methods

### Ex situ Experiment

The *ex situ* and *in situ* experiments took place at the apiary of the Chiang Mai University Meliponini and Apini Research Laboratory (MARL) in July 2025. Adult *T. mercedesae* mites were freshly obtained from highly infested *A. mellifera ligustica* colonies. Frames containing sealed brood were immediately transported to a nearby laboratory, where mite collection commenced without delay. Sealed brood cells were opened using epilation wax sheets. Mites were then collected individually from the comb using an insect sipper and transferred into a 50 mL Falcon tube. Subsequently, to preselect for vitality, chitinized adult individuals willing to actively climb to a syringe needle (18G) were used in the experiments one by one. The remaining mites in the vial were kept at 25°C for a maximum of 60 min to prevent a decrease in vitality; after this period, the stock was discarded, and fresh mites were collected.

To determine the effectiveness of lithium chloride over time, we adapted an assay previously developed for other mite species (Kolics et al., 2023). This method was optimized specifically for *Tropilaelaps* mites, as initial trials indicated that vortexing caused excessive mechanical stress to this species. For the assay, each selected mite was carefully placed onto a clean paper board strip.

Lithium chloride (LiCl) solutions were prepared using distilled water from powdered LiCl (Union Science Ltd., Chiang Mai, Thailand) to create the test concentrations ranging from 10.78 mM to 5.52 M (specifically: 10.78, 21.55, 43.11, 86.22 mM, and 0.17, 0.34, 0.69, 1.38, 2.76, 5.52 M). The exposure method involved precisely dropping the prepared solution directly onto each mite using a syringe needle (18G) of the same type used for mite selection. A fresh needle was used for each concentration tested to prevent cross-contamination. A 10-second exposure period was ensured by placing the entire drop onto the mite. For each concentration, an average of 20 mites were initially targeted, while 40 control individuals were exposed to distilled water.

We monitored for signs of lithium poisoning, which included the onset of tremorous movements, followed by the loss of ability to change location (cited as „Loc 1” hereafter), reduced locomotion with only slow leg movements (cited as „Loc stage 2” hereafter), and ultimately, the onset of death characterized by a complete absence of movement. Mites were monitored continuously, and the maximum observation period reached 17 hours post-exposure, under ambient laboratory conditions.

### In Situ Experiment

To demonstrate that the acaricidal effect of lithium could be elicited in situ as well, a pilot experiment was conducted. Four *A. m. ligustica* colonies were acquired from local beekeepers and were confined to Langstroth boxes all containing 6 frames. Three colonies were treated, and one served as an untreated control. Treated colonies were administered the treatment using the trickling method with 500 mM lithiated sugar syrup (1:1 water:sugar), as this was found to be the most effective concentration and way of administration used for *Varroa destructor* ^20^. Lithiated sugar syrup was trickled every third day, dripping 2.5 mL into the bee space of the standard Langstroth comb. Altogether, 8 treatments were applied to cover a whole drone brood cycle (24 days). Mite fall was counted on sticky boards coated with petroleum jelly (Vaseline) 3 days after each treatment. For experimental design see Table 1.

**Table 1.** Schedule of the *in situ* assay and treatment details for the treated and control colonies.

|  |  |
| --- | --- |
| 8<sup>th</sup> July | 1<sup>st</sup> treatment: 500 mM lithiated sugar syrup, control: sugar syrup) |
| 11<sup>th</sup> July | mite count, 2<sup>nd</sup> treatment (500 mM lithiated sugar syrup), control: sugar syrup) |
| 14<sup>th</sup> July | mite count, 3<sup>rd</sup> treatment (500 mM lithiated sugar syrup), control: sugar syrup) |
| 17<sup>th</sup> July | mite count, 4<sup>th</sup> treatment (500 mM lithiated sugar syrup), control: sugar syrup) |
| 20<sup>th</sup> July | mite count, 5<sup>th</sup> treatment (500 mM lithiated sugar syrup), control: sugar syrup) |
| 23<sup>th</sup> July | mite count, 6<sup>th</sup> treatment (500 mM lithiated sugar syrup), control: sugar syrup) |
| 26<sup>th</sup> July | mite count, 7<sup>th</sup> treatment (500 mM lithiated sugar syrup), control: sugar syrup) |
| 29<sup>th</sup> July | mite count, 8<sup>th</sup> treatments; lithium treated: formic acid, control: 500 mM lithiated sugar syrup |
| 1<sup>st</sup> Aug | mite count |

Concurrently for symptom analysis, we used modified sticky boards. Only the perimeter of the boards was coated in a 2 cm wide strip, leaving the central sticky area intact. This modification was done to prevent any mites falling alive from becoming stuck. Six hours after the first treatment, we collected mites falling over a 1 hour period from both the treated and control colonies. This was done to specifically observe the presence of the lithium-induced tremor symptom among the fallen mites. In total, 13 mites were obtained from the three treated colonies and 6 mites from the single untreated control.

### Statistical analysis

The sample sizes varied, with 18 animals observed for concentrations of 5 mM and 170mM, 17 for 86 mM, 340 mM, 690 mM, 1380 mM, and 5520 mM, 16 for 43 mM, 15 for 22 mM and 2760 mM, 13 for 11 mM and 31 for the control. Extreme values were identified, and 1 case (for concentration 690 mM) was excluded from further analysis because it exceeded 3 times the interquartile range.

A classic logistic growth curve (also known as the Pearl-Reed logistic curve) of the form *Y* = *K*/(1 + *b* . *e*^− *cX*^) was fitted to the mortality rate data for each treatment, of the form with *K =* 100, and *y* representing the mortality rate and *x* the exposure time. Parameters c and b define the position and slope of the fast-growth section of the curve, with *x = lnb /c* leading to *y = K/2*. This means that higher c values make the curve steeper, while with higher *b* values, the 50% value (*LC*_*50*_ stage) is reached later at higher *x* values.

The elapsed times from the beginning of the treatments were analyzed separately for each treatment for the recorded events detailed above.

The data were transformed using the natural logarithm *(ln (x+1))* and tested for normality using the Jarque–Bera and Shapiro–Wilk tests. The ln-transformed data for each stage were found to be normally distributed (p > 0.05).

The Levene test was used to justify homogeneous variances, and the Welch and Brown– Forsythe tests were applied to test for significant differences between the natural logarithms of exposure times to each stage (first tremorous movement and death) since the assumption of homogeneity of variances was violated.

Pairwise differences were analyzed with Tamhane’s post hoc test because of inhomogeneous variances.

The statistical tests were computed by SPSS 29.0 software (IBM, New York, NY, USA).

## Results

Our *ex situ* experiments revealed that lithium chloride is toxic to *T-. mercedesae* mites in a dose-dependent manner. We fitted classic logistic growth curves to the mortality rate data for each treatment concentrations (Figure 1). All models were significant with significant parameters (p < 0.001). The parameter values and R^2^ values indicating good fits of the estimated trend lines are shown in Table 2. As the concentration of LiCl increased, the time required to achieve 50% (*LC*_*50*_) and 90% (*LC*_*90*_) mortality decreased significantly (Figure 1 and Figure 2, Table 3).

**Table 2.** Parameters of the logistic growth curve models fitted to the *Tropilaelaps mercedesae* mortality data observed *ex situ* at different lithium chloride (LiCl) concentrations. The table details the sample size (n), the calculated model parameters (b and c), the coefficient of determination (R2) as a measure of fit, and the estimated time (in minutes) to reach 50% (*LC*_*50*_) and 90% (*LC*_90_) mortality for each concentration.

| Concentration<br>(M) | n<br>(Model) | Model* parameters |  | R<sup>2</sup> | Estimated time to |  |
| --- | --- | --- | --- | --- | --- | --- |
|  |  | b* | c* |  | *LC*<sub>50</sub> | *LC*<sub>90</sub> |
| 5520 | 17 | 4,30292 | -0,02354 | 0,84800 | 62,00 | 155,35 |
| 2760 | 15 | 7,77210 | -0,01490 | 0,93682 | 137,64 | 285,13 |
| 1380 | 17 | 11,62979 | -0,01856 | 0,96535 | 132,19 | 250,56 |
| 690 | 16 | 7,54541 | -0,01298 | 0,77782 | 155,75 | 325,09 |
| 340 | 17 | 10,94717 | -0,01369 | 0,93517 | 174,81 | 335,32 |
| 170 | 18 | 6,75250 | -0,01759 | 0,69538 | 108,59 | 233,51 |
| 86 | 17 | 23,34345 | -0,01372 | 0,88593 | 229,56 | 389,67 |
| 43 | 16 | 5,63254 | -0,00685 | 0,87190 | 252,47 | 573,38 |
| 22 | 15 | 10,85744 | -0,00831 | 0,91464 | 286,86 | 551,15 |
| 11 | 9 | 15,87059 | -0,00576 | 0,92695 | 480,36 | 862,15 |
| 5 | 15 | 18,02969 | -0,00331 | 0,88276 | 873,67 | 1537,45 |
| 0 | 16 | 53,60572 | -0,00250 | 0,88656 | 1590,58 | 2468,31 |

\*Indicates significant model with significant parameters at 0.001 significance level.

**Table 3.** Robust statistical analysis (Welch and Brown–Forsythe tests) of the effect of lithium chloride concentration on the time (ln-transformed data) required to reach toxicological stages (tremor, locomotion loss, death) in *T. mercedesae*, including effect size (*Eta* squared).

| Stages |  | Tremor | Loc 1 | Loc 2 | Death |
| --- | --- | --- | --- | --- | --- |
| Levene's Test Based on Mean | Statistic | 3.681 | 3.539 | 3.699 | 4.947 |
|  | Sig. | 0.000 | 0.000 | 0.000 | 0.000 |
| Welch Test | Statistic | 105.131 | 53.382 | 28.767 | 30.003 |
|  | Sig. | 0.000 | 0.000 | 0.000 | 0.000 |
| Brown–Forsythe Test | Statistic | 78.887 | 28.307 | 23.464 | 20.081 |
|  | Sig. | 0.000 | 0.000 | 0.000 | 0.000 |
| Eta Squared |  | 0,833 | 0,630 | 0,592 | 0,554 |

**Figure 1.**
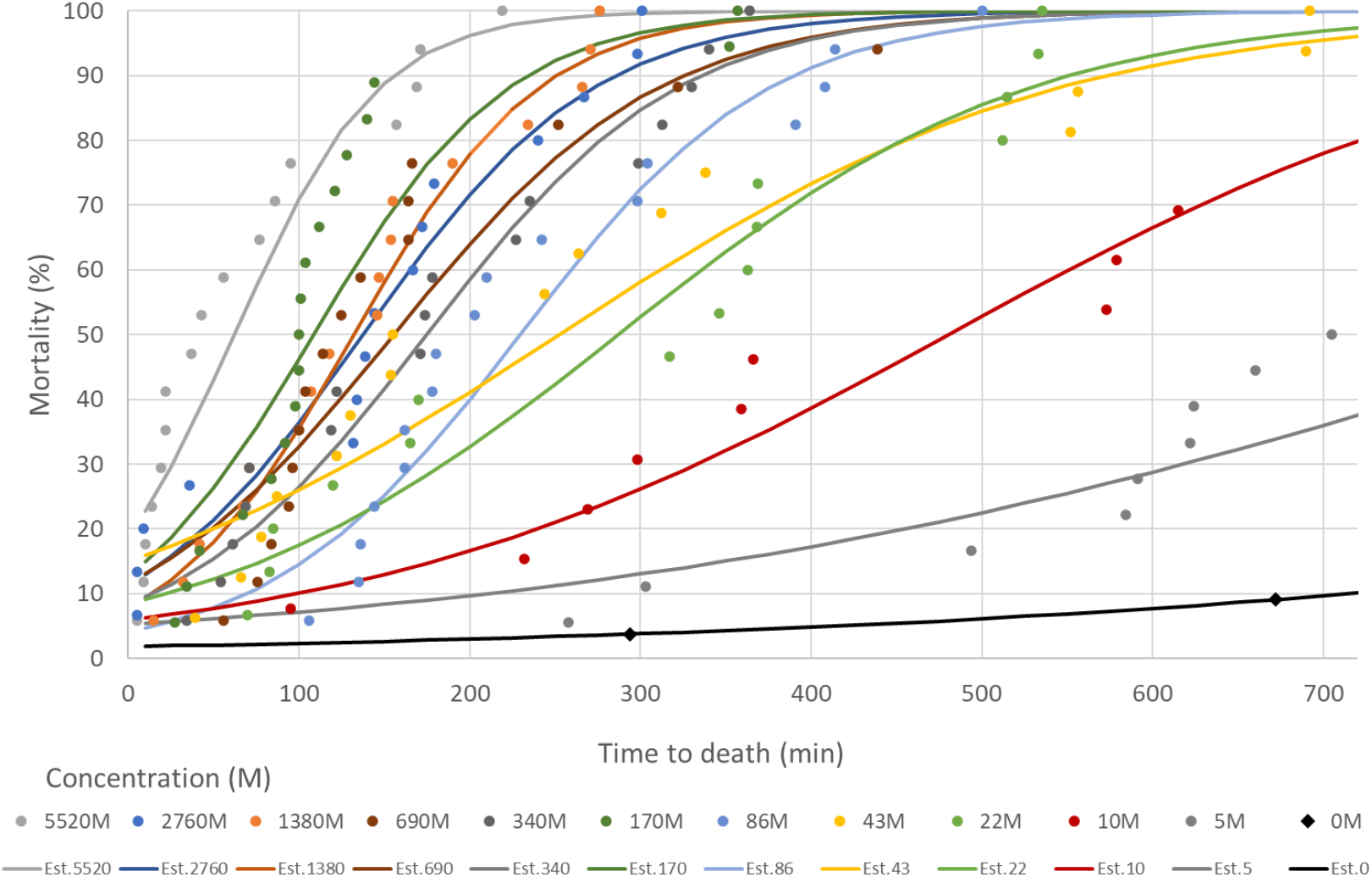
Mortality of *Tropilaelaps mercedesae* mites over time at different concentrations of lithium chloride. Each colored line represents the estimated mortality curve for a specific concentration, showing that higher concentrations lead to faster and higher mortality rates.

**Figure 2.**
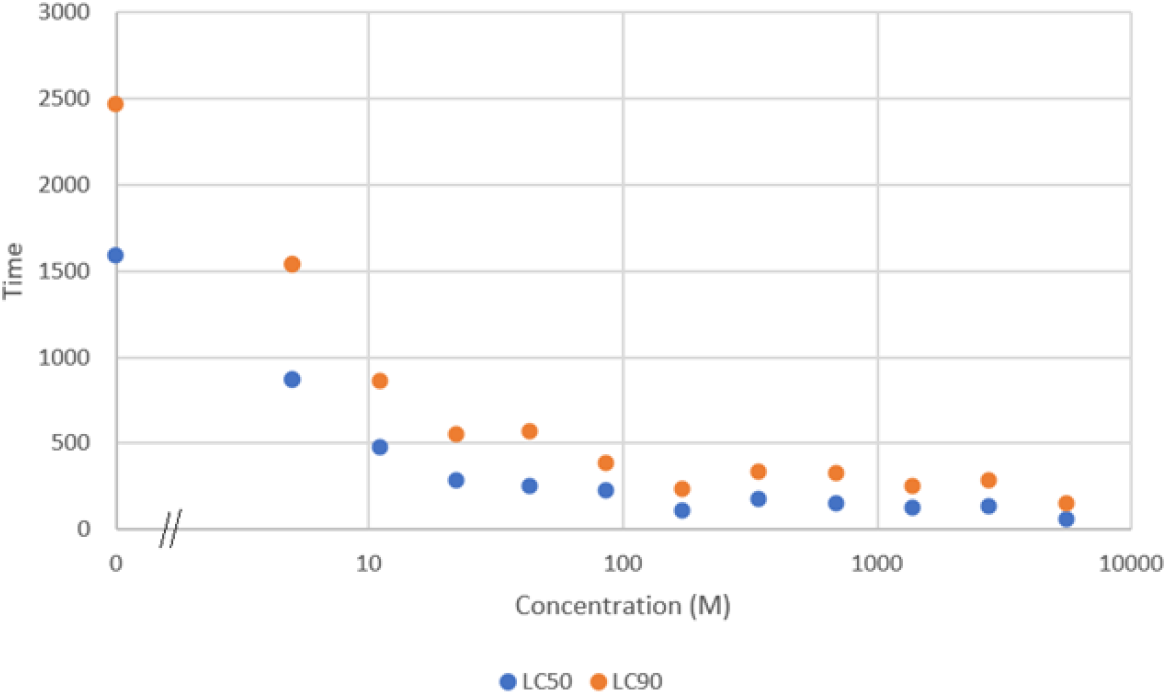
Estimated time (min) to reach *LC*_*50*_ (lethal concentration for 50% of mites) and *LC*_90_ (lethal concentration for 90% of mites) at various lithium chloride concentrations. The logarithmic scale for concentration highlights that the time to mortality decreases sharply as the concentration increases.

The onset of lithium poisoning symptoms, such as tremors, also showed a clear dose-dependent relationship. The observed mean time (natural logarithm transformed data, in minutes) elapsed after treatment needed to reach the given mortality stages for each treatment are shown Figure 3 and Figure 4.

**Figure 3.**
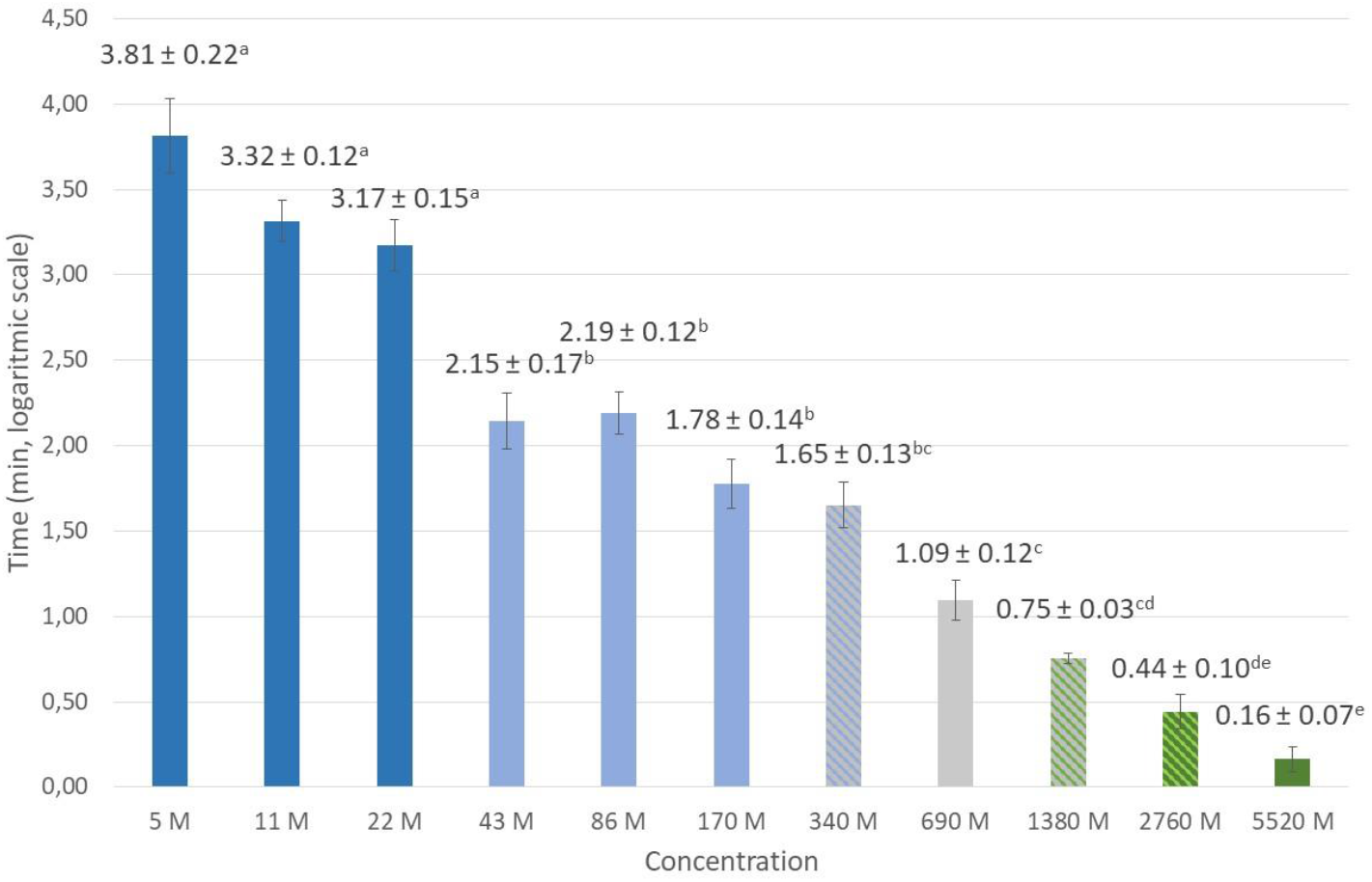
Mean time ± SE (standard error) on a logarithmic scale until the onset of tremor symptoms in *Tropilaelaps mercedesae* after exposure to different concentrations of lithium chloride. The letters (a, b, c, d, e) indicate statistically significant differences between the concentration groups, with lower letters corresponding to faster onset of tremors at higher concentrations. Error bars represent the standard error (SE).

**Figure 4.**
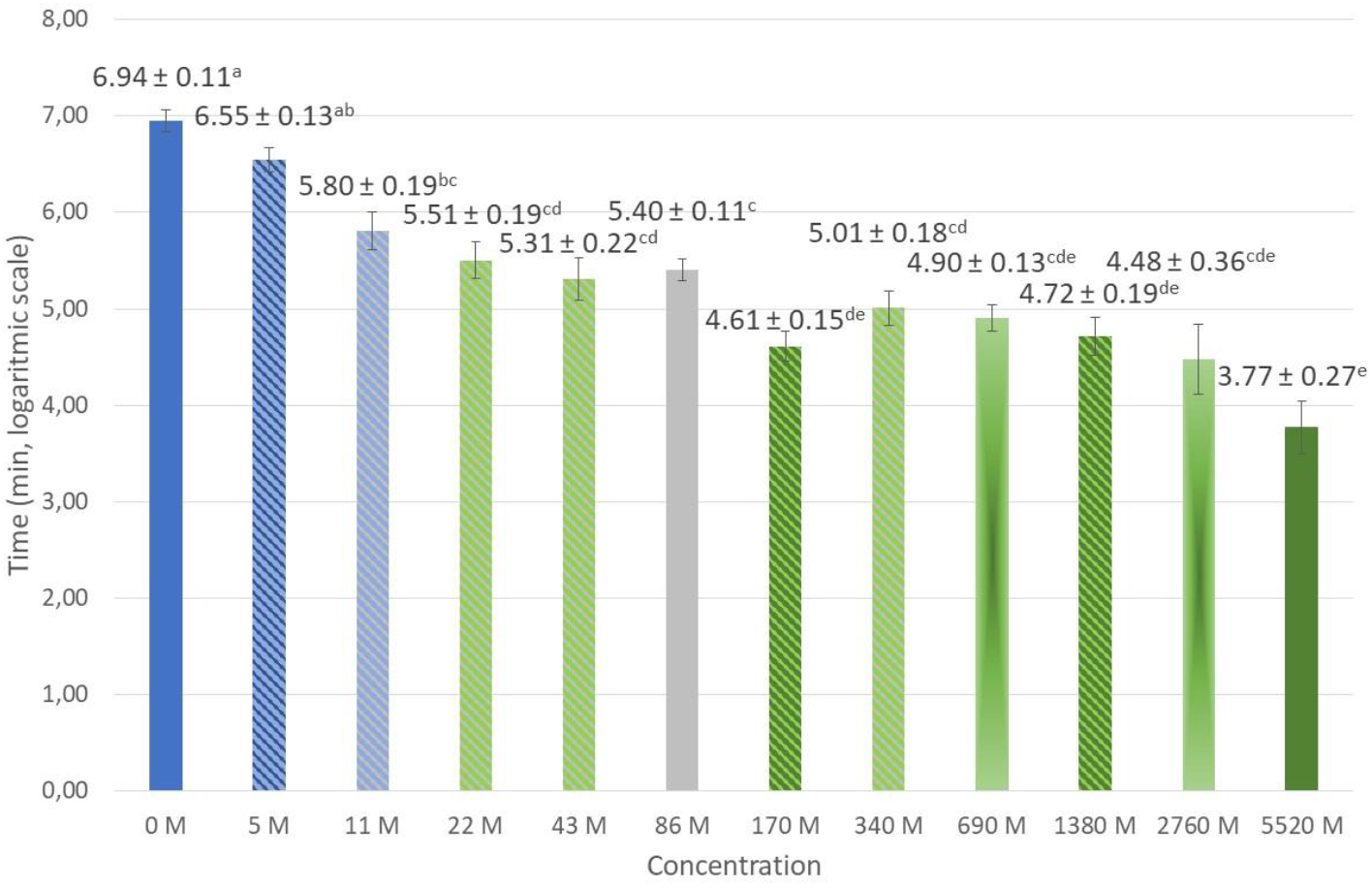
Mean time (on a logarithmic scale) until mortality stages in *Tropilaelaps mercedesae* following treatment with various concentrations of lithium chloride. The letters (a, b, c, d, e) denote significant differences between the groups, showing that higher concentrations lead to a quicker time of death. Error bars represent the standard error (SE).

Laboratory assays confirmed the high sensitivity of the species. The calculated 12-hour *LC*_*50*_ for *T. mercedesae* was 45.9 mM (Figure 5). The resulting toxicity profile demonstrates a notably higher intrinsic sensitivity in *Tropilaelaps mercedesae* compared to *Varroa destructor*, being more than 3.5 times more susceptible to lithium chloride.

**Figure 5.**
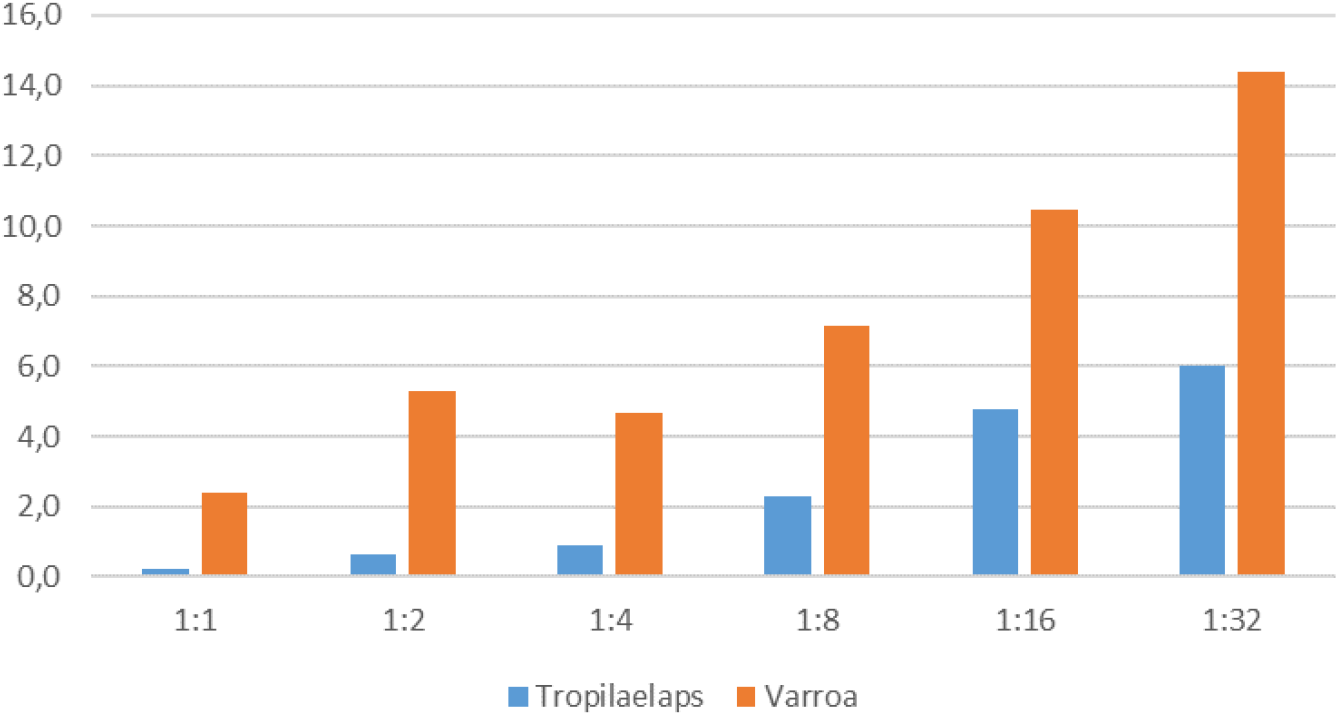
Treatment to tremor time (min) Comparative susceptibility of *Tropilaelaps mercedesae* and *Varroa destructor* to lithium chloride, expressed as 12-hour median lethal concentrations (*LC*_*50*_). *Tropilaelaps mercedesae* exhibits significantly higher sensitivity compared to *Varroa destructor*.

Statistical analysis confirmed significant differences (p < 0.001) observed between concentration groups for the onset of tremors and mortality, as well as for Loc 1 and Loc 2, as indicated by Welch, and Brown-Forsythe tests (Table 3). Consistent with this finding, higher concentrations of lithium chloride led to a significantly faster appearance of these tremors (Figure 3). Ultimately, the time until death was also significantly shorter at higher concentrations (Figure 4). The eta squared (*η*^*2*^) measures show the effect (concentration) size for analysis of variance. The value of eta square is 0.833 for the onset of tremors, and 0.554 for the mortality (Table 3).

In the *in situ* experiment, we monitored the dynamics of *Tropilaelaps* mite fall in three treated colonies and one untreated control colony (Table 4). The treated colonies (n=3) exhibited a generally decreasing trend in mite fall ver the 18-day, seven-application period. In contrast, the untreated control colony (n=1) showed a stable to slightly increasing background mite fall during the same period (e.g., 62 mites on day 3 vs. 86 on day 18). Crucially, direct evidence of the treatment’s *in situ* effect was observed when the control colony received its first lithium chloride application on day 21. This intervention caused an immediate and sharp increase in mite fall, rising from its 86-mite baseline (day 18) to 120 mites (day 21) and 149 mites (day 24) (Figure 6). This demonstrates that the elevated mite fall was a direct consequence of the lithium chloride application rather than other environmental factors.

**Figure 6.**
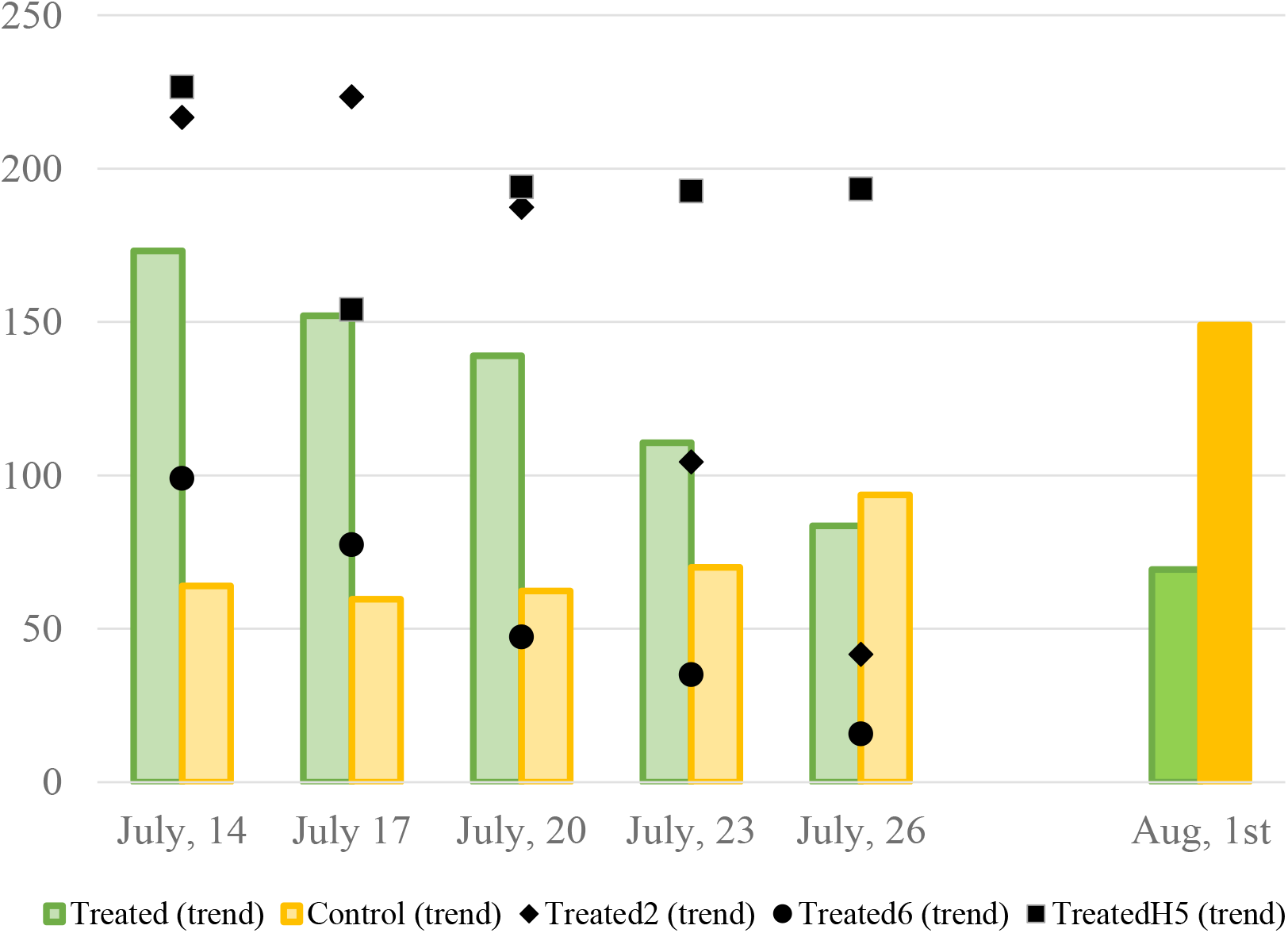
Temporal dynamics of *Tropilaelaps mercedesae* mite fall in lithium chloride-treated (n=3) and untreated control (n=1) colonies. The experimental timeline illustrates the primary treatment phase followed by a cross-over intervention on Day 21, where the administration of 500 mM lithium chloride to the control colony precipitated a marked increase in mite fall. The treated group received a terminal formic acid treatment.

**Table 4.** Cumulative *Tropilaelaps mercedesae* and *Varroa destructor* mite fall collected in three-day intervals from *in situ* treated (n=3; 500 mM lithium chloride trickling) and untreated control (n=1) colonies.*No data was collected due to broken sticky board.

This quantitative finding was supported by qualitative analysis. Symptoms were observed on living mites that had fallen onto modified (non-sticky) diagnostic bottom boards, used specifically for this purpose. Specifically, 8 out of 13 mites (61.5%) collected from the treated colonies exhibited these symptoms. In contrast, none of the mites (n=6) collected from the control colony exhibited these characteristic tremors associated with lithium poisoning, as identified in our *ex situ* assays. This supports that the observed mite fall was a result of the lithium-chloride treatment and not due to other environmental factors.

Taken together, these data demonstrate that the acaricidal potential of lithium chloride observed *ex situ* manifests *in situ* within a honey bee colony.

## Discussion

The expansion of *Tropilaelaps mercedesae* mites into temperate regions raises significant ecological and economic concerns for global apiculture. Crucially, recent evidence provided by Brandorf and colleagues challenges the prevailing assumption that this species is obligately dependent on continous presence of brood for survival ^8^. The potential for *T. mercedesae* to outcompete *V*.*destructor* in co-infested colonies, becoming the dominant biotic stressor, necessitates control strategies specifically adapted to its biology, which remains incompletely understood.

The impact of *Tropilaelaps* infestation may exceed even that of *V. destructor*, posing an imminent threat to European beekeeping and, by extension, agricultural pollination services. Furthermore, given the risk of acaricide resistance—mirroring the history of *V. destructor* management—there is a critical need to develop novel, effective therapeutic agents.

Our *ex situ* bioassays demonstrated the dose-dependent efficacy of lithium-chloride. Notably, the onset of tremors exhibited a significantly more refined dose-response relationship compared to mortality. This is evidenced by the greater number of statistically distinct treatment groups for tremor onset (Figure 3) relative to mortality (Figure 4). Furthermore, the substantially higher effect size (η^2^) for tremors (0.833) compared to death (0.554) quantitatively confirms that variations in concentration produce a more statistically distinguishable effect on this sublethal neurological symptom. The η^2^ values for the transitional stages, Loc 1 (0.630) and Loc 2 (0.592) follow this trend, indicating that the precision of toxicity assessment diminishes as physiological symptoms progress toward the terminal endpoint. Interestingly, the time of death –the most commonly used toxicity endpoint – yielded the lowest explanatory power (η^2^). These findings suggests that the characteristic tremor is not merely a precursor to death but serves as a more precise and immediate biomarker of the compound’s bioactivity. Consequently, we propose the onset of tremors as a valuable, potentially superior endpoint for comparative acaricidal assays. This metric may be particularly advantageous in studies comparing inter-specific sensitivity or in scenarios where determining the precise moment of death is complicated by the mite’s small size, slow toxicological progression, rapid mortality.

To contextualise our *ex situ* findings, lithium has demonstrated acaricidal activity against a diverse range of arthropods since its initial discovery in *Varroa destructor*. These include the tick *Dermacentor reticulatus* ^21^, the poultry red mite *Dermanyssus gallinae* ^22^, and the two-spotted spider mite *Tetranychus urticae* ^23^. Strikingly, *T. mercedesae* appears to be the most susceptible species identified to date. This exceptional sensitivity highlights lithium-chloride as a uniquely promising candidate for future *Tropilaelaps* management strategies.

When considering lithium residues, it is crucial to distinguish this mineral from conventional synthetic acaricides. Unlike xenobiotic toxins, lithium is a natural trace element often considered essential for physiological function ^24, 25^. Its biological impact is characterized by hormesis: while high concentrations can be toxic, necessitating strict dosage control ^26^, low doses - such as those potentially appearing in honey—are associated with positive physiological indications ^27^. Notably, micro-doses have been shown to provide neuroprotective benefits, including the stabilization of cognitive decline in Alzheimer’s disease ^28-30^. Therefore, the residue challenge here is not about zero tolerance, but about maintaining levels within the safe, beneficial range. Positively, similarly to organic acids ^31^, lithium is considered a natural component of honey ^17^. Our findings and previous studies suggest that lithium trickling treatments result in residues well below safety thresholds, presenting a manageable risk profile distinct from lipophilic synthetic compounds ^16, 32^.

Our in situ findings confirm that the acaricidal effect can be successfully elicited in honey bee colonies, even in the presence of brood. This is evidenced by the induced mite fall and the characteristic tremor symptoms, mirroring the neurological effects observed in all other lithium-exposed acari species examined to date ^21-23^. This consistency aligns with the established toxicological profile of lithium, as tremors constitute a primary clinical symptom of intoxication in vertebrates, including humans ^33^.

To assess practical relevance for apiculture, our pilot study demonstrated that acaricidal efficacy can be successfully achieved *in situ*. By employing the trickling method and concentration (500 mM) previously validated for *Varroa destructor* ^20^, we observed a clear anti-*Tropilaelaps* effect. Furthermore, the observation that surviving mites from the treated group exhibited tremor provides direct evidence linking the lithium treatment to the observed mite fall. This confirms that the recorded mortality was a direct consequence of the active substance rather than environmental factors. The potential of lithium is particularly noteworthy, as this mineral element may establish a unique therapeutic class for *Tropilaelaps* control, fundamentally distinct from conventional synthetic compounds, organic acids, or essential oils. This study provides the first toxicological evidence that *T. mercedesae* is highly sensitive to lithium chloride, exhibiting significantly lower *LC*_*50*_ values compared to *V*.*destructor*. We successfully translated these laboratory findings to colony conditions, demonstrating that the acaricidal effect - manifested by characteristic tremors and increased mite fall can be elicited *in situ*. While these pilot results validate lithium as a promising candidate with a unique mode of action, large-scale field trials are now required to quantify its practical efficacy and optimize application protocols for sustainable apicultural use.

## Conclusions

Our study establishes lithium chloride as a potent acaricide against Tropilaelaps mercedesae, demonstrating a sensitivity superior to that of Varroa destructor. The observation of characteristic tremors serves as a reliable biomarker for lithium intoxication, confirming that the treatment acts directly on the mites under field conditions. Unlike synthetic acaricides prone to resistance, lithium offers a unique mineral-based therapeutic perspective, potentially utilizing a mode of action distinct from current chemical treatments. Since lithium is a natural trace element in honey, it presents a promising “soft” chemical alternative. However, to fully integrate this novel agent into honey bee health programs, future research must focus on optimizing dosage regimes to maximize efficacy while maintaining residue levels within safe and beneficial ranges.

## Acknowledgments

We thank the Meliponini and Apini Research Laboratory (MARL) for assistance with colony maintenance and technical help.

## Funding

This work was supported by Pannonia Grant Program. The research was partially supported by Chiang Mai University and Kolics Apiaries.

## Notes

### Competing Interest Statement

The authors have declared no competing interest.

## Bibliograpy

1. Diagne C, Leroy B, Vaissière A-C, Gozlan RE, Roiz D, Jarić I, et al., High and rising economic costs of biological invasions worldwide. Nature 592: 571–576 (2021).

2. Traynor KS, Mondet F, de Miranda JR, Techer M, Kowallik V, Oddie MA, et al., Varroa destructor: A complex parasite, crippling honey bees worldwide. Trends in parasitology 36: 592–606 (2020).

3. de Guzman LI, Williams GR, Khongphinitbunjong K and Chantawannakul P, Ecology, life history, and management of Tropilaelaps mites. Journal of economic entomology 110: 319–332 (2017).

4. Woo K, Lee, JH The studies on the mites inhabiting the bee-hives. Korean Journal of Apiculture 8:140–156: 140–156 (1993).

5. Tokach R, Aurell D, Chuttong B and Williams GR, Observation of Tropilaelaps mercedesae (Mesostigmata: Laelapidae) on Western honey bees (Apis mellifera) exiting colonies. Journal of Economic Entomology 118: 966–969 (2025).

6. Franco S, Laurent, Marion, Duquesne, Véronique, Geographical Spread of the Exotic Mite Tropilaelaps spp.: State of Play of the Worldwide Situation in March 2025. Scientific Note published on the website of the European Union Reference Laboratory for Bee Health 2025).

7. Nabian s, akhzari s, Ahmadi K, Garami Sadeghian A and Ali A, A review of the reemerge mite Tropilaelaps Clareae in Iran. Honeybee Science Journal 15: 51–60 (2024).

8. Brandorf A, Ivoilova MM, Yañez O, Neumann P and Soroker V, First report of established mite populations, Tropilaelaps mercedesae, in Europe. Journal of Apicultural research 64: 842–844 (2025).

9. Janashia I, Uzunov A, Chen C, Costa C and Cilia G, First report on Tropilaelaps mercedesae presence in Georgia: The mite is heading westward! Journal of Apicultural Science 68: 183–188 (2024).

10. Tokach R, Chuttong B, Aurell D, Panyaraksa L and Williams GR, Managing the parasitic honey bee mite Tropilaelaps mercedesae through combined cultural and chemical control methods. Scientific Reports 14: 25677 (2024).

11. Chaimanee V, Warrit N, Boonmee T and Pettis JS, Acaricidal activity of essential oils for the control of honeybee (Apis mellifera) mites Tropilaelaps mercedesae under laboratory and colony conditions. Apidologie 52: 561–575 (2021).

12. Ziegelmann B, Abele E, Hannus S, Beitzinger M, Berg S and Rosenkranz P, Lithium chloride effectively kills the honey bee parasite Varroa destructor by a systemic mode of action. Scientific reports 8: 683 (2018).

13. Garbian Y, Maori E, Kalev H, Shafir S and Sela I, Bidirectional transfer of RNAi between honey bee and Varroa destructor: Varroa gene silencing reduces Varroa population. PLoS pathogens 8: e1003035 (2012).

14. Kolics É, Mátyás K, Taller J, Specziár A and Kolics B, Contact Effect Contribution to the High Efficiency of Lithium Chloride Against the Mite Parasite of the Honey Bee. Insects 11: 333 (2020).

15. Prešern J, Kur U, Bubnič J and Šala M, Lithium contamination of honeybee products and its accumulation in brood as a consequence of anti-varroa treatment. Food chemistry: 127334 (2020).

16. Kolics É, Sajtos Z, Mátyás K, Szepesi K, Solti I, Németh G, et al., Changes in Lithium Levels in Bees and Their Products Following Anti-Varroa Treatment. Insects 12: 579 (2021).

17. Bogdanov S, Jurendic T, Sieber R and Gallmann P, Honey for nutrition and health: a review. Journal of the American College of Nutrition 27: 677–689 (2008).

18. Hernández O, Fraga J, Jiménez A, Jiménez F and Arias J, Characterization of honey from the Canary Islands: determination of the mineral content by atomic absorption spectrophotometry. Food chemistry 93: 449–458 (2005).

19. Kolics É, Specziár A, Taller J, Mátyás KK and Kolics B, Lithium chloride outperformed oxalic acid sublimation in a preliminary experiment for Varroa mite control in pre-wintering honey bee colonies. Acta Veterinaria Hungarica 68 2021).

20. Kolics B, Kolics É, Mátyás K, Taller J and Specziár A, Comparison of Alternative Application Methods for Anti-Varroa Lithium Chloride Treatments. Insects 13: 633 (2022).

21. Kolics B, Mátyás K, Solti I, Bacsi Z, Kovács S, Specziár A, et al., Efficacy of In Vitro Lithium Chloride Treatments on Dermacentor reticulatus. Insects 14: 110 (2023).

22. Kolics B, Kolics É, Solti I, Bacsi Z, Taller J, Specziár A, et al., Lithium Chloride Shows Effectiveness Against The Poultry Red Mite (Dermanyssus gallinae). Insects 13: 1005 (2022).

23. Solti I, Kolics É, Keszthelyi S, Bacsi Z, Staszny Á, Nagy E, et al., Evaluation of the Acaricidal Activity of Lithium Chloride against Tetranychus urticae (Acari: Tetranychidae). Horticulturae 8: 1127 (2022).

24. Pickett EE and O’Dell BL, Evidence for dietary essentiality of lithium in the rat. Biological trace element research 34: 299–319 (1992).

25. Schrauzer GN, Lithium: occurrence, dietary intakes, nutritional essentiality. Journal of the American College of Nutrition 21: 14–21 (2002).

26. McKnight RF, Adida M, Budge K, Stockton S, Goodwin GM and Geddes JR, Lithium toxicity profile: a systematic review and meta-analysis. The Lancet 379: 721–728 (2012).

27. Calabrese EJ, Pressman P, Hayes AW, Dhawan G, Kapoor R, Agathokleous E, et al., Lithium and hormesis: Enhancement of adaptive responses and biological performance via hormetic mechanisms. Journal of Trace Elements in Medicine and Biology 78: 127156 (2023).

28. Nunes PV, Forlenza OV and Gattaz WF, Lithium and risk for Alzheimer’s disease in elderly patients with bipolar disorder. The British Journal of Psychiatry 190: 359–360 (2007).

29. Damiano RF, Loureiro JC, Pais MV, Pereira RF, Corradi MdM, Di Santi T, et al., Revisiting global cognitive and functional state 13 years after a clinical trial of lithium for mild cognitive impairment. Brazilian Journal of Psychiatry 45: 46–49 (2023).

30. Aron L, Ngian ZK, Qiu C, Choi J, Liang M, Drake DM, et al., Lithium deficiency and the onset of Alzheimer’s disease. Nature 2025).

31. Bogdanov S and Martin P, Honey authenticity: a review. Mitt Lebensm Hyg 93: 232–254 (2002).

32. Prešern J, Kur U, Bubnič J and Šala M, Lithium contamination of honeybee products and its accumulation in brood as a consequence of anti-varroa treatment. Food chemistry 330: 127334 (2020).

33. Amdisen A, Clinical and serum-level monitoring in lithium therapy and lithium intoxication. Journal of Analytical Toxicology 2: 193–202 (1978).

